# Integrative single-cell multi-omics of CD19-CAR^pos^ and CAR^neg^ T cells suggest drivers of immunotherapy response in B-cell neoplasias

**DOI:** 10.1101/2024.01.23.576878

**Authors:** Mercedes Guerrero-Murillo, Aina Rill-Hinarejos, Juan L. Trincado, Alex Bataller, Valentín Ortiz-Maldonado, Daniel Benitez-Ribas, Marta Español, Europa Azucena González, Nuria Martinez-Cibrian, Doménica Marchese, Lourdes Martín-Martín, Alejandro Martin Garcia-Sancho, Holger Heyn, Manel Juan, Álvaro Urbano-Ispizúa, Julio Delgado, Alberto Orfao, Elisabetta Mereu, Clara Bueno, Pablo Menendez

**Author notes:** Correspondence should be addressed to: -Elisabetta Mereu -Clara Bueno -Pablo Menendez Josep Carreras Leukemia Research Institute. Barcelona. Spain. These authors equally contributed to this study (Mercedes Guerrero-Murillo, Aina Rill-Hinarejos, Juan L. Trincado).

## Abstract

How phenotypic, clonal, and functional heterogeneity of CAR-T-cells impact clinical outcomes remain understudied. Here, we integrated clonal kinetics with transcriptomic heterogeneity resolved by single-cell omics to explore cellular dynamics response of both non-transduced (CAR^neg^) and transduced (CAR^pos^)T-cells. CAR^neg^ and CAR^pos^T-cells were longitudinally interrogated in the manufactured infusion product (IP) and *in-vivo* at CAR-T cell expansion peak in five B-ALL patients treated with CD19CAR-T-cells (varni-cel). Significant differences were found in the cellular dynamics between CAR^pos^ and CAR^neg^T-cells in response to therapy. CAR^pos^T-cells in the IP exhibited a significant higher CD4:CD8 ratio than CAR^neg^T-cells, and the CD4:CD8 CAR^pos^T-cell composition impacted therapy outcome as confirmed in a larger cohort of 24 varni-cel-treated B-ALL patients. Conversely, an inverted trend in the CD4:CD8 CAR^pos^T-cell ratio was consistently observed at the expansion peak, with clonally expanding CD8^+^ effector memory and cytotoxic T-cells being the most abundant populations. Expanded cytotoxic CAR^pos^γδT cells emerged at the expansion peak, and the extent of their *in-vivo* expansion positively correlated with treatment efficacy, which was validated in a large cohort of B-ALL patients (n=18) treated with varni-cell and B-cell lymphoma patients (n=58) treated with either lisa-cel or axi-cel. Our data provide insights into the complexity and diversity of T-cell responses following CAR-T cell therapy and suggest drivers of immunotherapy response.

## Introduction

Chimeric antigen receptor (CAR)-T cell therapy has emerged as a groundbreaking treatment option for relapsed/refractory (r/r) hematological malignancies, particularly B-cell acute lymphoblastic leukemia (B-ALL), B-cell lymphoma and multiple myeloma, as shown by impressive clinical complete response (CR) rates after infusion of CD19-directed CAR-T cells (CD19CAR-Ts)^1–5^ for B-ALL and lymphoma, and BCMA-directed CAR-T cells^6^ for multiple myeloma. Unfortunately, sustained CR rates at one year following treatment have been observed in <50% of these patients, indicating a pressing need to better understand the underlying cellular and molecular features of CAR-T cell therapy leading to different patient outcomes.^7,8^

The therapeutic performance of CAR-T cells depends on multiple factors, including the design of the CAR construct, antigen density, the quality and starting phenotype of the T cells sourced for manufacturing, the lymphodepletion and infusion scheme, the tumor burden, and the tumor microenvironment.^9–14^ Additionally, clinico-biological factors that confer immune resistance to the tumor cell are key determinants of clinical response and durability, but identifying the factors that predict clinical outcomes remains challenging.

Recent advances in single-cell (sc) multi-omics approaches have provided new insights into the clonal kinetics and heterogeneity of CAR-T cell products, enabling the identification and characterization of CAR-T cell populations associated with sustained remission^15–21^. However, these studies have also described variability in the clonal composition and transcriptional signatures of CAR-T cells and how this translates into the distinct functional T-cell subsets, highlighting the need for additional studies to identify those T-cell subtypes/clones responsible for short-term efficacy and long-term persistence^12,10^.

Here, we integrated clonal kinetics using single-cell αβTCR sequencing (sc-αβTCR-seq) with transcriptomic heterogeneity resolved by single-cell RNA sequencing (scRNA-seq) to explore the cellular dynamics response of both non-transduced (CAR^neg^) and transduced (CAR^pos^) T cells sorted by flow cytometry based on CAR expression. We examined both (CAR^neg^ and CAR^pos^) T-cell fractions in the same manufactured infusion product (IP) and in peripheral blood (PB) at the time of CAR-T cell expansion peak following infusion (peak) in 5 adult r/r B-ALL patients treated with the CD19CAR-T product varnimcabtagene autoleucel (varni-cel) ^8,22,23^. Our data reveal significant differences in the cellular dynamics between CAR^pos^ and CAR^neg^ T cells in response to CD19CAR-T cell therapy. Expanded cytotoxic CAR^pos^ γδT cells, not traceable by αβTCR, emerged at the expansion peak in all patients, and the extent of *in vivo* expansion of CAR^pos^ γδT cells correlated with treatment efficacy. Clinic-biological associations were validated in larger cohorts of CD19CAR-T-treated B-ALL (varni-cel) and Diffuse Large B-cell Lymphoma (DLBCL; either lisa-cel or axi-cel) patients. This study highlights the complexity and diversity of the T cell response following CAR-T cell therapy, which may not solely depend on CAR^pos^ αβT cells.

## Results

### Multimodal identification of T cell subtypes in response to CD19CAR-T cell therapy

To investigate immune responses to CD19CAR-T cell therapy and understand the impact of CAR expression on T cell identities, we analyzed the single-cell transcriptional and immune repertoire profiles of transduced (CAR^pos^) and non-transduced (CAR^neg^) T cells in the IP and at peak expansion in 5 patients with r/r B-ALL who received the CD19CAR-T product varnimcabtagene autoleucel ^8,22,23^ (**Fig 1A**). Clinico-biological information of the patients from the discovery cohort is shown in **Table 1**. CAR-T cells were successfully manufactured from leukapheresis from all patients enrolled in the study. After 1:1 CD4:CD8 T cell selection, the cells were activated using anti-CD3/anti-CD28 beads and transduced with the CD19-specific CAR using a lentiviral vector. Following transduction, the cells were expanded *ex vivo* for 8 (+/-1) days, and 10^6^ CAR-T cells/kg body weight was infused into each patient in three sequential administrations (10:30:60)^24^. All patients received cyclophosphamide (900 mg/m2) and fludarabine (90 mg/m2) lymphodepleting chemotherapy prior to CAR-T cell infusion, as per protocol^22^. CAR-T cell transduction levels in the final IP ranged between 25% and 40%, and the infused CAR-T cells exhibited different degrees of expansion in patients, with peak expansion occurring between week 1-4 post-infusion (**Fig 1B, Fig S1A**). Patient outcomes were monitored for up to 12 months, during which time 3 out of 5 patients experienced relapse (**Fig 1C**). In two patients (pt1 and pt4) who experienced short remission periods (2 and 3 months, respectively) both detectable CAR^pos^ T cells and leukemic cells coexisted.

**Figure 1.**
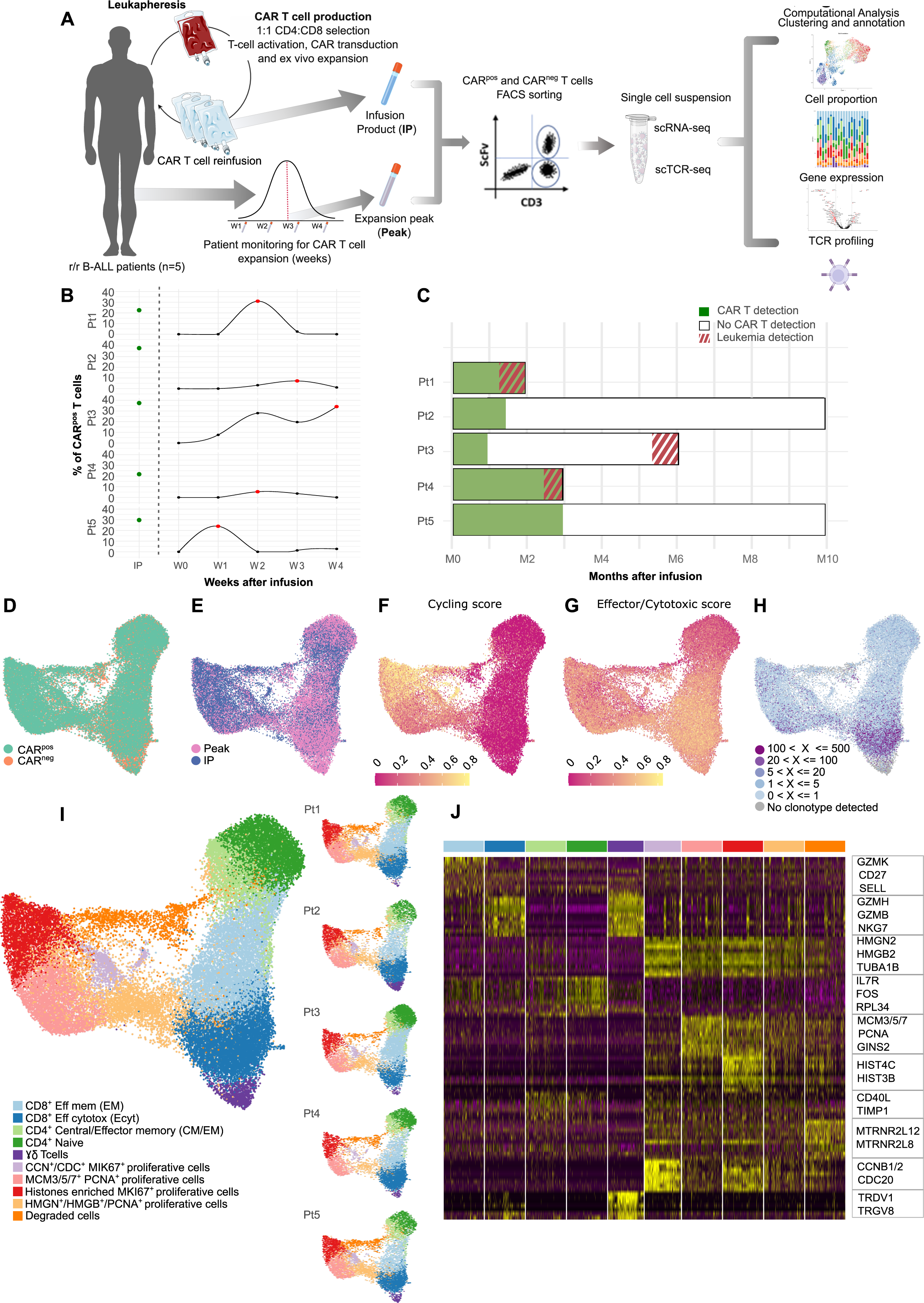
The landscape of T cell populations in r/r B-ALL patients treated with varnimcabtagene autoleucel. **A)** Schematic overview of the experimental design. CAR-T cells were manufactured from leukapheresis products. CD4/CD8 T cells were selected, activated, transduced with the CD19CAR construct, and expanded. Upon manufacturing, a sample was collected from each patient’s product prior to infusion, conforming the “Infusion Product” (IP). Subsequently, patients were monitored weekly to assess the CAR-T cell expansion, and the sample with the highest level of CAR^pos^ T cells over this period conformed the peak of expansion (Peak). CAR^neg^ and CAR^pos^ T cell fractions were FACS-purified from IP and Peak samples and prepared in single-cell suspensions for scRNAseq and scTCRseq. **B)** Flow cytometry monitoring for CAR^pos^ T cell expansion over the first four weeks after infusion. The percentage of CAR^pos^ T cells in the IP is indicated as a green dot whereas Peak samples (timepoints) are represented as red dots. **C)** Summary of the clinical evolution of the patients over a ten-month follow-up. Solid green and white bars indicate the presence or absence (no detection) of CAR^pos^ T cells in disease-free patients, respectively. Stripped red lines indicate patient relapse. **D–G)** Uniform manifold approximation and projection (UMAP) of the 38,190 cells pooled from all the samples from the 5 patients colored as follows: (**D**) CAR^pos^ *vs* CAR^neg^ cells, (**E**) IP *vs* Peak, (**F,G**) scaled from high (yellow) to low expression (magenta) of cycling score (**F**), and effector/cytotoxic score (**G**). **H)** UMAP representing clonotype sizes (from grey to purple). **I)** UMAP colored by the 10 different cell clusters identified by unsupervised clustering both combined (left) and separated by patient (right). **J)** Heatmap of the top 50 cluster-specific gene markers identified by differential expression analysis.

**Table 1.**
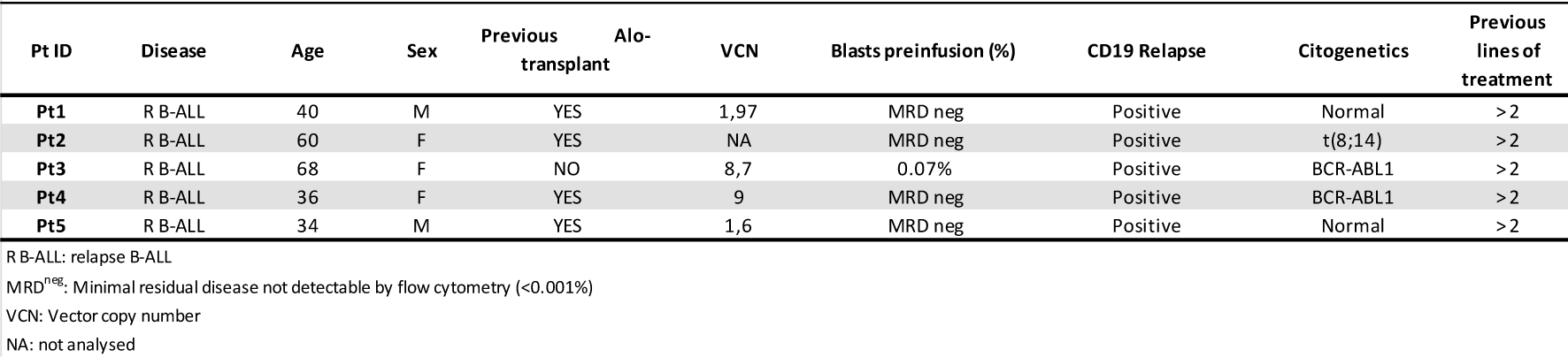
Clinical and biological features of the patients included in this study.

In total, we analyzed 38,190 high-quality T cells from all samples and time points. The number of sequenced cells was comparable between the experimental conditions (**Table S1**). CAR transcripts were evenly distributed in all patients in both IP and peak samples; they were only minimally detected in CAR^neg^ cells of the corresponding samples and were filtered out (**Fig S1B,C, Table S2**). To identify distinct T cell subpopulations and characterize their transcriptional heterogeneity, we first normalized and scaled the single-cell gene expression counts, and we next performed principal component analysis (PCA) on the highly variable genes to reduce the dimensionality of the data and maintain only the most biologically informative genes. Then, we used a shared nearest-neighbor graph-based clustering method coupled with uniform manifold approximation and projection (UMAP) to identify and visualize in 2D cells with similar transcriptional profiles (see Methods). There was a high overlap between cells from the CAR^pos^ and CAR^neg^ subpopulations (**Fig 1D**), while T cells tended to be more distinct when based on their sampling time (**Fig 1E**). Specifically, the peak samples exhibited a higher abundance of non-cycling clonally expanded effector/cytotoxic T cells when compared with the IP samples (**Fig 1E-H**). The analysis of cluster-specific gene expression using those genes differentially expressed (DEGs) among clusters allowed us to identify 10 subtypes of T cells, encompassing a wide range of differentiation states, from the less mature and proliferating T cells to the most differentiated CD8^+^ effector memory and cytotoxic T cells (**Fig 1I,J, Fig S1D,E**).

Four clusters of highly proliferative cells were identified based on the expression of genes involved in cell cycling, DNA replication, and proliferation, including cyclins/cyclin-dependent kinases, HMGB proteins, *PCNA*, *MCM5*, *MCM7*, *MKI67*, and *AURKA* (**Fig 1I,J**). These clusters also exhibited a high cycling score (**Fig 1F**). Additionally, a CD4^+^ naïve T-cell cluster was identified that expressed *IL7R*, with low cycling and effector cytotoxic scores. Central effector memory markers were expressed in the remaining clusters, including one CD4^+^ and two CD8^+^ T-cell clusters (**Fig 1I,J**). One CD8^+^ cluster contained effector cytotoxic markers, including *GZMH*, *GZMB*, *GNLY*, *NKG7*, and *PRF1*, whereas the other TCD8^+^ cluster expressed *GZMK*, indicative of a different T cell population with an effector memory profile, as previously described^17,25^. Of note, a separate cluster of cytotoxic γδT cells expressing *GNLY*, *NKG7*, *GZMH*, and *GZMB* as well as the γδ markers TCR delta variable 1 (*TRDV1*) and TCR gamma variable 8 (*TRGV8*), but lacking αβTCRs, was identified (**Fig 1G-J**). Lastly, a cluster with high expression of mitochondrial and ribosomal genes was identified and classified as degraded cells. While all T cell subtypes were present in each patient, the Morisita index revealed no shared TCR sequences across patients, indicating a lack of similarity between their TCR repertoires. However, we observed some overlap in the TCR repertoires between the CAR^pos^ and CAR^neg^ subpopulations within the same patient (**Fig 1H, Fig S1F**). Overall, these findings provide a comprehensive characterization of the different T cell populations and their molecular profiles, which can aid in understanding the CD19CAR-T cell response.

### Changes in subtype composition between CAR^pos^ and CAR^neg^ T cells in the IP link to treatment outcome

To understand the impact of CAR transduction during manufacturing, we compared T cell subtype compositions between CAR^pos^ and CAR^neg^ T cells at the IP (**Fig 2A,B**). Analysis revealed that proliferation was the primary source of variation upon manufacturing, with higher number of proliferative cells found within the CAR^pos^ T cell population (**Fig 2C**). However, when examining proliferative and non-proliferative cells separately, we observed a consistent cell subtype composition of proliferative cells in all patients. By contrast, a significant shift in the abundance of CD8^+^ T cells was observed in non-proliferating cells, with CAR^neg^ T cells showing a significantly higher proportion of CD8^+^ effector cytotoxic and memory T cells (**Fig 2C,D**).

**Figure 2.**
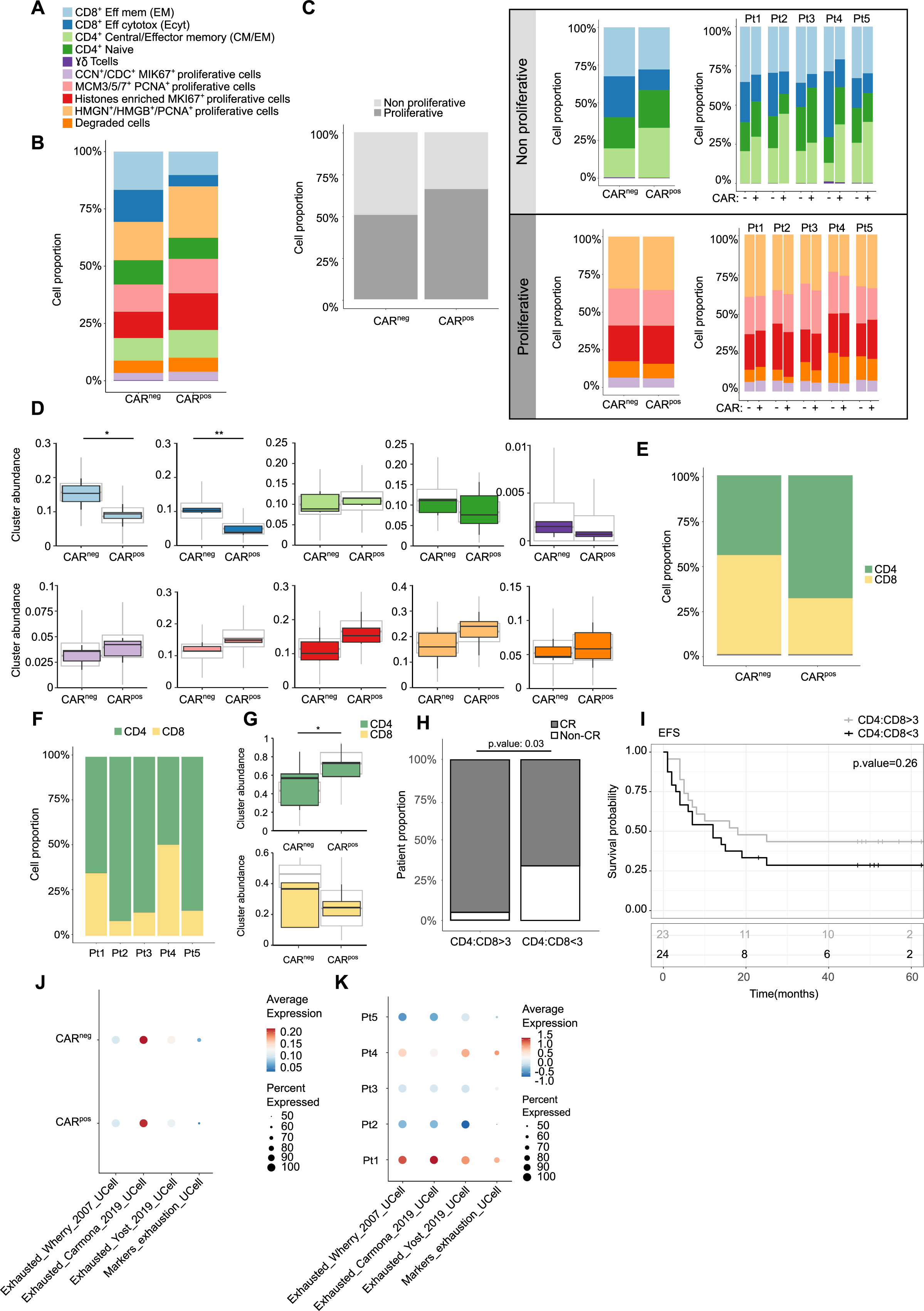
Differences among CAR^pos^ and CAR^neg^ T cells in the IP. **A)** Color code used to identify the different T cell populations. **B)** Stacked bar plot showing T cell subtype composition of CAR^pos^ and CAR^neg^ T cells in the IP. **C)** Left panel, proportion of proliferative and non-proliferative CAR^pos^ and CAR^neg^ T cells in the IP. Stacked bar plots showing the T cell subtype composition within the non-proliferative (right upper panel) and proliferative (right lower panel) CAR^pos^ and CAR^neg^ T cells both overall and by patient. **D)** Boxplots with the differential abundance analysis of CAR^pos^ and CAR^neg^ T cell clusters. The expected abundance is indicated in grey boxes and the observed abundance colored by T cell subtype. **E,F)** Composition of the predicted CD4 and CD8 T cell subtypes both overall (**E**) and by patient (**F**). CD4/CD8 T cells were identified by a well-established reference-based classifier matchsore2 using a published dataset as a reference and Azimuth (see Methods). **G)** Boxplots with the differential abundance analysis of CAR^pos^ and CAR^neg^ CD4 and CD8 T cell populations in the IP. **H)** CR rates achieved in a retrospective cohort (n=47) of patients with r/r B-ALL treated with varnimcabtagene autoleucel in whom CAR-T cell products had a CAR^pos^ CD4:CD8 ratio > 3 or <3 (p.value=0.03). **I)** Kaplan-Meier survival curve for EFS. Time is represented in months. Grey and black lines represent patients with a CAR^pos^ CD4:CD8 ratio >3 and <3, respectively. **J,K)** Dot plots showing the expression of four exhaustion signatures in CAR^pos^ and CAR^neg^ T cells both overall (**J**) and by patient (**K**). The Ucell method was used to score each signature (Table 3).

To gain further insight into the composition of the T cell populations and potential functional implications following CAR transduction, we assigned a CD4 or CD8 phenotype to each cell by projecting the cells onto different publicly available single-cell multi-omic references obtained from 24 peripheral blood mononuclear cell (PBMC) samples that were processed using CITE-seq and scRNA-seq and the human CD8^+^ and CD4^+^ TIL atlas (see Methods) (**Fig S2A**). We found a consistent and substantial shift in the abundance of CD4^+^ T cells in CAR^pos^ products across all patients (**Fig 2E-G**). This higher representation of CD4^+^ T cells in the CAR^pos^ fraction of the IP was independent of CD4 T cell-specific activation or tonic signaling (**Fig S2B,C**). Of note, a diminished proportion of CAR^pos^ CD4^+^ T cells in the IP was associated with very early relapses (**Fig 1C, 2F**). This was further validated by flow cytometry in a retrospective cohort (n=47) of varmi-cel-treated r/r B-ALL patients (**Table S3**). In this expanded cohort, 95% and 60% of patients infused with a CAR^pos^ CD4:CD8 ratio >3 or <3, respectively, achieved CR (p-value=0.03) (**Fig 2H**). Accordingly, five-year event-free survival (EFS) was superior in patients who had received a CAR T cell product with a CAR^pos^ CD4:CD8 ratio >3 (43.5% [IC 27.3%–69.3%] vs 28.5% [IC 15%–54.3%]) (**Fig2H,I, S2D**). Together, FACS analysis in the validation cohort confirmed the scRNAseq data in the discovery cohort to suggest the CD4:CD8 CAR^pos^ T cell composition as a predictor of CAR-T cell therapy outcomes.

Additionally, we assessed relevant transcriptional signatures related to T cell activation and exhaustion and tonic signaling that might affect the clinical outcome of the CAR-T cell therapy (**Table S4**) and found no significant differences between CAR^pos^ and CAR^neg^ T cell populations in the IP (**Fig 2J, S2B,C**). However, patients with very early relapse had consistently higher scores of different exhaustion signatures, irrespective of the expression of the CAR construct **(Fig 2K, S2E, Table S4**), indicating a CAR-independent exhausted T cell response in these patients, which could potentially be used as biomarkers to predict patient outcomes following CAR-T cell therapy.

### Transcriptomic and clonal dynamics of CAR^pos^ T cells from *ex vivo* manufacturing (IP) to *in vivo* expansion peak

To gain a more comprehensive understanding of the CAR-T cell response dynamics, we investigated the evolution of CAR^pos^ T cells at both the transcriptional and clonal levels. By comparing changes in T cell composition from the IP to the expansion peak in the patient, we identified a significant decline in proliferative cells, which were subsequently replaced by CD8 T cells (**Fig S3A,B and Fig 3A left,**). Within non-proliferative cells, an inverted trend in the CD4:CD8 ratio was observed at the expansion peak in all patients, with CD8 T cell effector memory and cytotoxic subtypes representing 75% of the cellular product **(Fig 3A right, S3B**). Of note, a population of CAR^pos^ γδT cells emerged at the expansion peak in all patients (**Fig 3A**).

**Figure 3.**
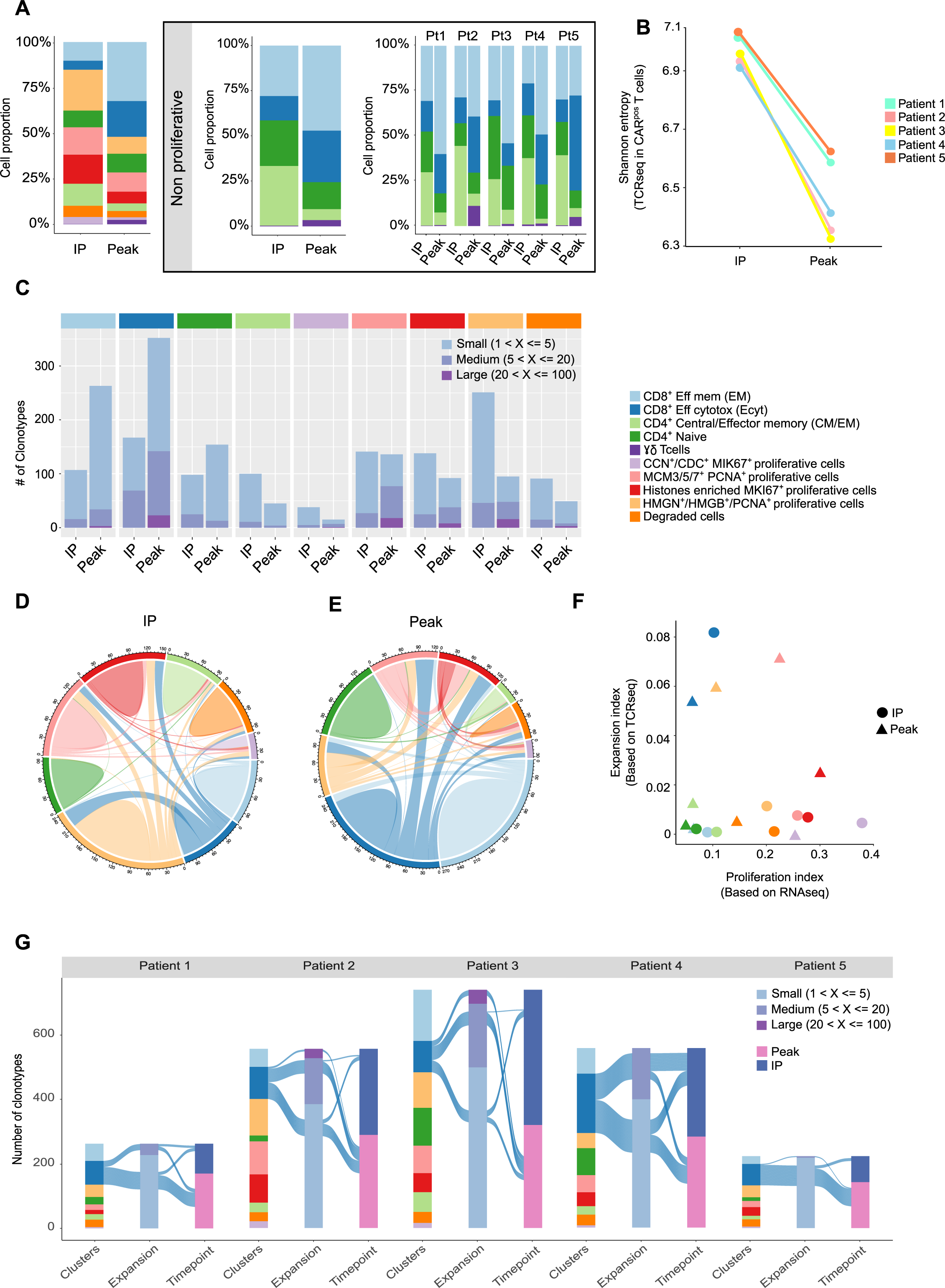
Longitudinal analysis of CAR^pos^ T cells across timepoints (IP and Peak). **A)** Left panel, stacked barplots of CAR^pos^ T cell subtype composition in IP and Peak. Right panel, stacked barplots of CAR^pos^ T cell subtype composition within the non-proliferative T cells in IP and Peak both overall and by patient. **B)** IP-Peak paired Shannon entropy measure calculated for each patient to evaluate the diversity of the TCR repertoire within the CAR^pos^ T cells. **C)** Occupied TCR repertoire space for CAR^pos^ T cells by cell type/cluster and timepoint (IP and Peak). Only small (1<x<=5), medium (5<x<=20), and large (20<x<=100) clonotypes were considered. **D,E)** Chord diagrams showing the interconnection of shared expanded clonotypes of the different T cell subtypes/clusters in IP (**D**) and Peak (**E**). **F)** Scatterplot showing the clonal expansion and the proliferation indexes of the CAR^pos^ T cells at IP (circles) and Peak (triangles). Colors indicate the T cell cluster as shown in Fig 2A. Proliferation index was calculated based on scRNAseq data, and expansion index was computed by Startrac software from scTCRseq data (see Methods). **G)** Alluvial plot indicating the total number of cells sharing clonotype between IP and Peak per patient. Each bar is colored to show the proportion of the different T cell subtypes (clusters), expansion and timepoint samples per patient. Blue lines connect the CD8^+^ effector memory T cells clonotypes with their information.

To better explore the complexity of the CAR-T cell responses and quantify the diversity of TCR repertoires, we employed the Shannon entropy index, which is a diversity measure that considers both abundance and distribution of TCR sequences. Comparison of the Shannon entropy indices between IP and peak revealed a consistent trend indicative of an *in vivo* clonal expansion of CAR^pos^ T cell clones over time for all the patients (**Fig 3B**). Indices of evenness and richness including the ACE and the inverse Pielou index were also consistent with these results (**Fig S3C**). This expansion of CAR^pos^ T cell clones was particularly pronounced within the CD8 T cell fraction, with the CD8 cytotoxic subtype showing the largest number of clonotypes and magnitude of expansion (**Fig 3C, 1H**).

Notably, the process of clonal expansion and diversification of the CD8 cytotoxic cells seemed to be continuous, occurring from the IP to the peak (**Fig S3D**). As the process unfolded, the emergence of shared clones between CD8 cytotoxic cells and other proliferative T cells increased at the expansion peak (**Fig 3D,E**), indicating the ongoing diversification of the T cell response. It is possible that the high expansion of CD8^+^ T cells provides a larger pool of cells available for differentiation and recruitment into other T cell subsets. This is supported by the finding that CAR^pos^ CD8^+^ cytotoxic T cells have a high expansion score but a low proliferative score (**Fig 3F**), suggesting that their increase in number is not due to cell division but instead to cell survival, recruitment, or differentiation. By contrast, CAR^pos^ CD4^+^ T cell subtypes exhibited the lowest expansion and proliferation scores, indicating that CAR-T cell responses are characterized by the preferential expansion and survival of specific T cell subsets, rather than the global expansion of all T cell populations.

We further investigated the T cell response following CAR-T cell therapy by comparing the expansion and clonal diversity between patients with varying CAR-T cell responses. This analysis revealed notable differences in T cell expansion and clonal diversity between patients (**Fig S3E,F**). We observed a consistent expansion of CD8 cytotoxic T cells in all patients at the peak (**Fig S3F**). However, while we observed some shared T cell clones between the IP and peak in all patients, the number of shared clones and their rate of expansion did not correlate with treatment response (**Fig 3G**). Overall, our findings suggest that the expansion of CD8 cytotoxic T cells may play an important role in determining the response to CAR-T cell therapy, but other factors may also be involved in determining the overall outcome of the treatment. In addition, the difference in the patterns of clonal expansion and sharing between patients may reflect differences in the underlying mechanisms of CAR-T cell response and the complexity of the T cell repertoire.

### *In vivo* expansion of CAR^pos^ γδT cells links to a more favorable treatment outcome

We initially identified a population of CAR^pos^ T cells characterized by high cytotoxic activity (**Fig 1G**), a high number of non-classifiable CD4 or CD8 T cells (**Fig S2A**), and undetectable αβTCR expression (**Fig 1H, 4A**). We determined that these cells are cytotoxic γδT cells based on their expression of the γδ TCR markers *TRDV1* and *TRDV8* and cytotoxic genes, such as granzymes, *PRF1*, *GNLY*, and *NKG7* (**Fig 4B,C**). The absolute number of these CAR^pos^ γδT cells was very low upon manufacture (IP) but was highly expanded *in vivo* (between 2-100-fold) at the expansion peak (**Fig 4D**). Of note, the two patients with a massive *in vivo* expansion of CAR^pos^ γδT cells (pt2 and pt5) had a better clinical outcome in the discovery cohort (**Fig 1C, 4D**). This was further validated by flow cytometry in two retrospective larger cohorts of CD19CAR-T cell treated r/r B-ALL and DLBCL patients (**Table S3**). In the B-ALL setting, the number of varni-cel-treated patients who displayed *in vivo* expansion of CAR^pos^ γδT cells was ∼3-fold higher among disease-free patients than in relapsing patients one year after treatment (**Fig 4E**, 28% *vs* 10%, n=18 patients). This finding was also confirmed in an independent cohort of r/r DLBCL patients (n=58) treated with axi-cel (n=33) and tisa-cel (n=25) among whom CAR^pos^ γδT cell counts ≥5 cells/ml at the expansion peak was associated with a significantly much better progression/recurrence-free survival (p<0.001, **Fig 4F**). Collectively, these results suggest that CAR^pos^ γδT cells may play a potential role in treatment efficacy. We were, however, unable to further investigate this expansion using TCR sequencing due to the unique nature of the γδ TCR.

**Figure 4.**
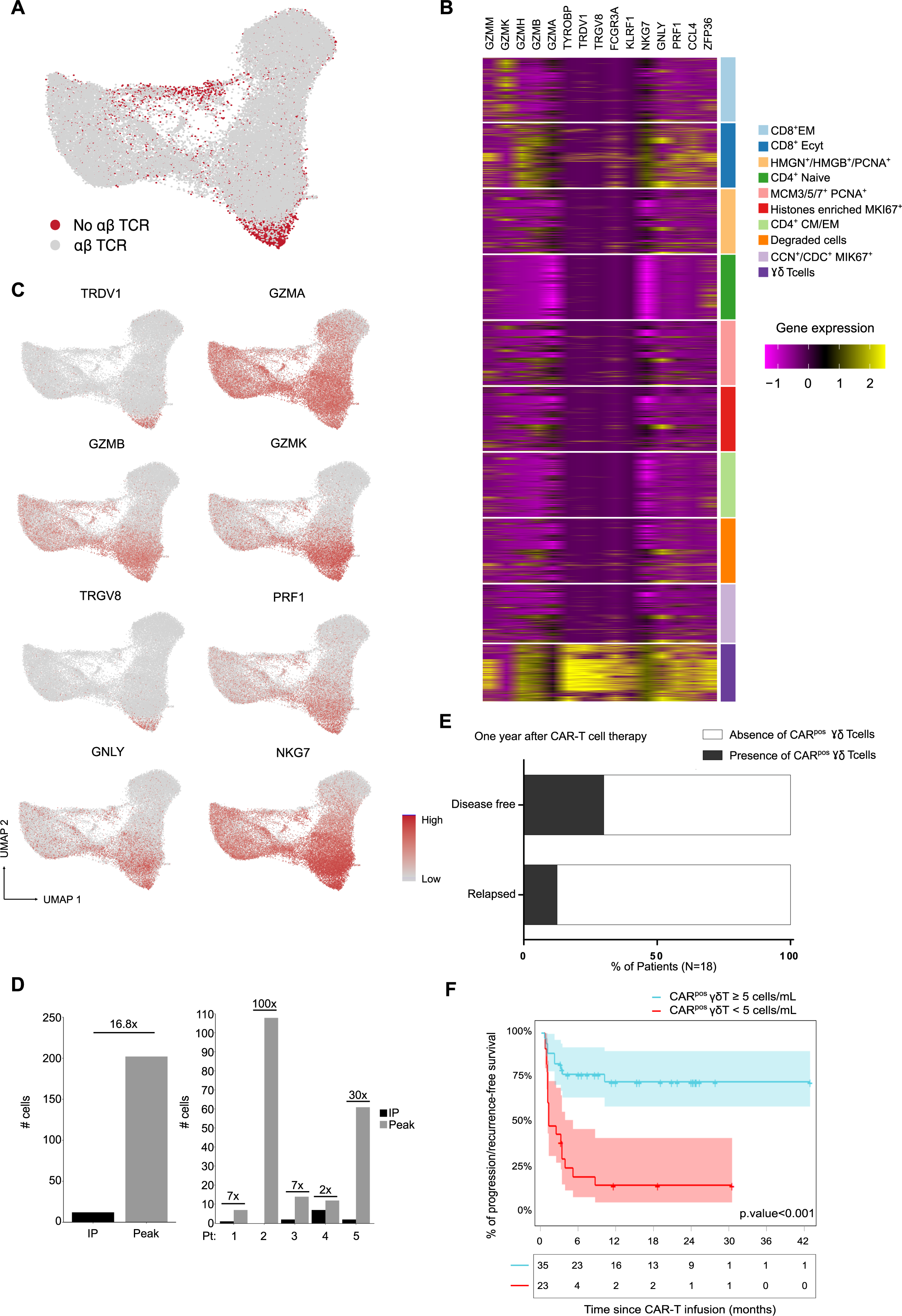
*In vivo* expansion of CAR^pos^ γδT cells correlates with treatment outcome. **A)** UMAP plot colored with the presence (grey) or absence (red) of TCR αβ sequences. **B)** Heatmap of the top 15 gene markers of γδT cells identified by differential expression analysis and cytotoxic genes. **C)** UMAP plots colored by the expression of the indicated γδ and cytotoxicity genes. **D)** Bar plots showing the absolute number of the identified γδT cells at IP (black) and Peak (grey) combining all samples (left panel) and by patient (right panel). **E)** Proportion of disease-free and relapsed B-ALL patients one year after treatment with varni-cel who displayed an *in vivo* expansion of CAR^pos^ γδT cells. The presence of CAR^pos^ γδT cells was assessed by flow cytometry in a retrospective/historical cohort (n=18). **F)** Prognostic impact of the number of CAR^pos^ γδT cells in PB at the expansion peak on progression/recurrence-free survival of r/r 58 DLBCL patients treated with axi-cel and lisa-cel products.

## Discussion

CAR-T cell therapy has emerged as an effective and safe therapeutic approach in patients with r/r hematological malignances. Our biological understanding and clinical management of such an adoptive immunotherapy are still in their early stages, but undoubtedly, it is poised to become a therapeutic intervention during earlier stages of the disease. CAR-T cell therapy will eventually benefit many more patients and even a broader spectrum of tumors when significant logistical and biological milestones are achieved. In turn, a significant reduction of manufacturing times and costs is needed. On the other hand, reliable predictive biomarkers of clinical success, measured as the persistence of CAR-T cells capable of long-term disease control without relapse or immune escape, are in high demand. In this regard, the design of CAR constructs, the target of choice, antigen density, the quality and phenotype of the manufactured T cells, the lymphodepletion and infusion scheme, the tumor burden, the tumor microenvironment, and other factors that confer immune resistance to the tumor cell are key determinants of clinical response and durability.

The identification of reliable markers that predict clinical outcomes in CAR-T-treated patients remains suboptimal. In an attempt to contribute to this, we have integrated single-cell clonal kinetics with transcriptomic heterogeneity to explore the cellular dynamics response of both non-transduced (CAR^neg^) and transduced (CAR^pos^) T-cell fractions longitudinally sampled from the IP and from PB at the time of CAR-T cell expansion peak^22^. Although recent advances in single-cell multi-omics have provided relevant information into the clonal kinetics and heterogeneity of CAR-T cell products ^15–21,26^, this is to the best of our knowledge the more complete study analyzing patient-matched CAR^pos^ and CAR^neg^ T fractions, without underestimating the potential impact of CAR^neg^ T cells in the clinical response. Our cohort included patients who did not respond to the treatment and relapsed early after infusion in the presence of detectable CAR-T cells, but also patients who evidenced a protracted CR and/or delayed relapse in the absence of CAR^pos^ T cells. Neither age nor sex, molecular/cytogenetic features of the disease, CAR-T expansion, disease burden or the number of CAR^pos^ T cells at infusion correlated with the clinical outcome in our discovery cohort^6,27–29^ suggesting either leukemic cell autonomous mechanisms of disease progression or an impact of the variability in the clonal, phenotypic and functional composition of the infused T cells in patient outcome.

Our results unravel significant differences in the cellular dynamics between CAR^pos^ and CAR^neg^ T cells and highlight the complexity and diversity of the T cell responses following CD19CAR-T cell therapy. Upon manufacturing, a consistent and substantial shift in the abundance of CD4^+^ T cells was observed in CAR^pos^ T-cell fraction across all patients, which was not associated with CD4 T cell-specific proliferation, activation or tonic signaling. Importantly, the CAR^pos^ CD4:CD8 ratio in the IP was linked to clinical outcome. A CAR^pos^ CD4:CD8 ratio <3 was associated with lower CR rates and lower EFS as validated by flow cytometry in a retrospective cohort of 47 patients with r/r B-ALL treated with varni-cel. Of note, although no differences in T-cell exhaustion, activation or tonic signaling signatures could be found between CAR^pos^ and CAR^neg^ T cells in the IP, those patients with very early relapse had consistently higher scores of different exhaustion signatures, irrespective of the expression of the CAR construct, indicating that a CAR-independent T cell dysfunction could potentially predict patient outcomes following CAR-T cell therapy. These findings are consistent with previous studies suggesting that the CD4:CD8 T cell ratio and exhaustion signatures are useful predictors of clinical response^16,30,31^. It might thus be possible to identify those patients with a higher risk for relapse or who might benefit from additional therapies to enhance the durability/persistence of the T-cell response. Additionally, these data provide clues about the potential advantage of pre-selecting less exhausted T cell populations, perhaps immediately after the leukapheresis procedure, before CAR-T cell product manufacturing.

The expression of the CAR serves as a label/identifier to trace the cellular dynamics, proliferation, expansion, and clonality of CAR-T cells from the IP to the peak of *in vivo* expansion in the patient. The CAR^neg^ T-cell fraction was not analyzed at the peak of *in vivo* expansion because it is not possible to unequivocally determine if these cells were initially present at the IP or, on the contrary, are T cells *de novo* generated in the patient during the 4 weeks that elapse from the infusion to CAR-T cell expansion. Integrated single-cell clonal kinetics with transcriptomic heterogeneity revealed a consistently higher abundance (75%) of non-cycling clonally expanded effector/memory cytotoxic CD8^+^ T cells in the peak samples. Of note, two separate cytotoxic CD8^+^ clusters expressing different granzymes and cytotoxic genes were observed, indicating the existence of distinct effector memory CD8^+^ T cells, as previously described^17,25^. Overall, although clonal expansion and TCR diversification of the CD8^+^ T cells seemed to be continuous from the IP to the peak, the emergence of shared clones between CD8 cytotoxic cells and other proliferative T cells increased at the expansion peak, highlighting the ongoing diversification of the T cell response. The fact that CAR^pos^ CD8^+^ cytotoxic T cells have a high expansion score, but a low proliferative score, suggests that the high expansion of CD8^+^ T cells may provide a larger pool of cells available for differentiation and recruitment into other T cell subsets, suggesting that their increase in number is not due to cell division but instead to cell survival, recruitment, or differentiation. By contrast, CAR^pos^ CD4^+^ T cell subtypes exhibited the lowest expansion and proliferation scores, indicating that a preferential expansion and survival of specific T cell subsets, rather than the global expansion of all T cell populations, underlie CAR-T cell responses.

Important, integrated scTCR/scRNA-seq data also uncovered an independent cluster of CAR^pos^ cytotoxic T cells expressing multiple GZM genes, *PRF1*, GNLY, and *NKG7* which could not be sequenced through standard αβTCR sequencing. These T-cells were identified as γδT cells based on their expression of γδT cell-specific genes, *TRDV1* and *TRDV8*. The absolute number of these CAR^pos^ γδT cells was very low at the IP but was highly expanded *in vivo* (up to 100-fold) at the expansion peak. Intriguingly, a massive *in vivo* expansion of CAR^pos^ γδT cells was associated to a better clinical outcome as validated by flow cytometry in retrospective cohorts of patients with r/r B-ALL treated with varni-cel and patients with r/r DLBCL treated with tisa-cel or axi-cel. This result was somewhat surprising given that i) γδT cells represent <5% of the total number of all T cells in PB, and ii) the manufacturing process of CAR-T cells includes selection of CD4 and CD8 cells while >70% of γδT cells are CD4^-^ and CD8^-32^. The relevance of γδT cells in CAR-T immunotherapies may be due not only to their high cytotoxic capacity but also to the intrinsic high capacity of γδT cells to migrate from blood and infiltrate the bone marrow and other peripheral tissues^33,34^. This highlights the complexity and diversity of the T cell response following CAR-T cell therapy, which may not solely depend on CAR^pos^ αβT cells, and suggest that future CAR-T cell therapy production could benefit from not filtering out the vast majority of CD4^-^CD8^-^ γδT cells (and potentially also other innate T and T/NK cells), which have previously been reported to play a crucial role in decade-long leukemia remissions after CD19CAR-T cell therapy^15^. Notably, an *in-silico* study of 18,000 patients with cancer established that a tumor-associated γδT cell gene signature also correlated significantly with improved overall survival across 39 different malignancies, further advocating the application of γδT cells in cancer therapy^35^. Further experimental and clinical studies are needed to prospectively assess the clinical impact of γδT cells in CAR-T-treated patients with B-ALL and other blood cancers.

Despite the limited number of patients in the discovery cohort, the robustness of our results is enhanced by the convergence of findings from both experimental investigations and clinical validations in varni-cel-treated B-ALL patients and lisa-cel/axi-cel-treated DLBCL patients, reinforcing the significant correlation between higher CAR^pos^ CD4:CD8 T cell ratio at IP and the higher *in vivo* expansion of CAR^pos^ γδT cells and clinical outcome.

## Materials and Methods

### Patient samples and sample collection

We analyzed T-cells from five adult r/r B-ALL patients treated with the CD19CAR-T-cell product varnimcabtagene autoleucel (varni-cel ^8,22,23^. Varni-cel was produced and infused as detailed elsewhere^24^. After CAR-T infusion, patients were followed up by FACS in both PB (weekly for the first 5 weeks post-infusion) and BM for disease recurrence and CAR-T-cell persistence as previously described^24^. Mononuclear cells (MNCs) were isolated by density gradient centrifugation from the manufactured IP (pre-infusion) and from PB the week of CAR-T cell expansion peak following infusion (peak). Detailed clinical and biological information of patients from the discovery cohort is summarized in **Table 1**. Additional 47 r/r B-ALL patients treated with varni-cel in the context of the original phase I trial, compassionate use, or hospital exemption ^8,22,23^, as well as 58 r/r DLBCL patients treated with axi-cel (n=33) and tisa-cel (n=25) were included in the expanded validation cohorts. The study was approved by the Institutional Review Boards of the Hospital Clinic of Barcelona (r/r B-ALL) and University Hospital of Salamanca (r/r DLBCL) (**Table S3**).

### FACS sorting

MNCs from either IP or peak were immunophenotyped by FACS using the monoclonal antibodies CD3-FITC (BD Biosciences, San Jose, CA), CD4-FITC (BD Biosciences), and CD8-PerCP Cy5.5 (BD Biosciences). The cell surface expression of the CD19CAR was traced by using AffiniPure F(ab’)₂ fragment goat anti-mouse IgG (H+L)-APC (Jackson ImmunoResearch, Westgrove, PA). CAR^pos^ and CAR^neg^ CD3+ T-cells were FACS-sorted at high purity (>95%) using a FACS Aria flow cytometer (BD Biosciences).

### Library preparation and scRNA-seq and sc-αβTCR-seq

CAR^pos^ and CAR^neg^ CD3+ T-cells were centrifuged for 5 minutes at 500×g at 4°C to a concentration of 700–1200 cells/μL. Cell concentration and viability were verified using the TC20™ Automated Cell Counter (Bio-Rad Laboratories, Hercules, CA) after staining the cells with Trypan blue. Cells were partitioned into Gel BeadInEmulsions (GEMs) with a target cell recovery of 5000 total cells per sample using the Chromium Controller system (10× Genomics, Pleasanton, CA). Gene expression (GEX) and TCR-enriched sequencing libraries were prepared using the Chromium Single Cell 5’ Library and Gel Bead Kit v1.0 (10× Genomics, #1000006) and the Chromium Single Cell V(D)J Enrichment Kit, Human T Cell (10× Genomics, #1000005). Briefly, after GEM-RT clean up, cDNA was amplified for 14 cycles and quantified with an Agilent Bioanalyzer High Sensitivity Chip (Agilent Technologies, Santa Clara, CA). GEX libraries were prepared using 1–25 ng of cDNA and indexed with 16 cycles of PCR using the PN-220103 Chromium i7 Sample Index Plate. For the preparation of αβTCR-enriched libraries, 11 cycles of TCR target enrichment PCR 1 and 2 were performed and the resulting cDNA was quantified on an Agilent Bioanalyzer High Sensitivity chip (Agilent Technologies). TCR libraries were prepared and indexed with 9 cycles of amplification using the PN-220103 Chromium i7 Sample Index Plate. The size distribution and concentration of the libraries were verified on an Agilent Bioanalyzer High Sensitivity chip (Agilent Technologies). Finally, sequencing of GEX and TCR libraries was carried out on an Illumina NovaSeq 6000 sequencer to obtain approximately 20,000 reads per cell and 5000 reads per cell, respectively.

### Processing of scRNA-seq and scTCR-seq

Cell Ranger software (10× Genomics; v3.1.0) was used to demultiplex the BCL files into FASTQ files using the *cellranger mkfastq* pipeline. The different FASTQ files were aligned to the reference genome (version hg38) with the addition of the CAR construct sequence using the *cellranger count* pipeline. The Cell Ranger Single-Cell Software Suite (v3.1.0) with VDJ option was used to map TCR-enriched libraries to the GRCh38 VDJ pre-built reference provided by 10× Genomics. Cell Ranger performed default filtering for quality control, and produced barcodes.tsv, genes.tsv, and matrix.mtx files for each sample. Clonotype data were included into the metadata of each sample. In total, 38,190 T cells and 31,863 productive TCR sequences were obtained (**Table S1**).

### scRNA-seq data analysis

The following downstream analyses were performed with Seurat library (v4.3.0)^36^ for R 4.0.5. Prior to data integration, we analyzed each sample individually and performed a quality control and filtering to remove possible apoptotic, damaged cells, or doublets as well-established. Filtered-out cells were identified as those with <200 genes detected, >20% expressed mitochondrial genes and log10(UMI counts)>3. In addition, a very limited (<1%) number of myeloid cells expressing *CD14* or *LYZ* were detected and excluded from the analyses. A few contaminating CAR^pos^ T cells found within the FACS-sorted CAR^neg^ T cell fraction were also excluded. A total of 38,190 T cells were sequenced with an average of 7,638 cells per patient. 19,531 and 18,659 of the sequenced T cells were CAR^pos^ and CAR^neg^, respectively. 19,016 and 19,174 of the sequenced T cells were from IP and peak, respectively (**Tables S1,S2**). After filtering, the *LogNormalize* method was used to normalize the expression data, and the 3000 most variable genes were obtained with *FindVariableFeatures*. For data integration, we selected 3000 genes with *SelectIntegrationFeatures* function and used them to scale and run the PCA on each sample. The selected genes were also used as input for the *FindIntegrationAnchors* function. A set of 2000 anchors were obtained where we excluded cell cycle genes and TRV genes (both *TRAV* and *TRBV*). All samples were finally integrated with *IntegrateData* using the set of anchors and the first 30 dimensions. The integrated dataset was scaled with *ScaleData* and regressing out S and G2M cell cycle markers to shift mean expression across cells to 0, following the integration Seurat pipeline. The 20 first principal components were used to perform the linear reduction after a PCA exploration. Then, cell clusters were obtained with *FindNeighbors* and *FindClusters*, with a resolution of 0.5 and visualized with the UMAP algorithm *RunUMAP*.

### CD4 and CD8 projection

The classification of CD4 and CD8 T cells based on direct gene expression analysis can be challenging due to the sparsity of scRNA-seq data. To address this issue, we utilized projection-based classification methods by leveraging previously published reference datasets. Specifically, we used Azimuth^36^ a software with a PBMC dataset^37^ comprising 11,769 cells that were sequenced by scRNA-seq and CITEseq. This approach allowed us to classify T cells with high sensitivity. Additionally, we employed the ProjecTILs software^38^ to transfer labels from two other reference datasets, the human CD8^+^ and CD4^+^ TIL atlas^39^, which contained between 11 and 12 thousand sequenced T cells. To ensure reliable classification, we required both Azimuth and ProjecTILs to agree on whether a cell was CD4 or CD8. Cells that were not classified or showed disagreement between the two software tools were excluded from the classification.

### Cluster analysis, cell type classification and signature analysis

The first 20 principal components with a resolution of 0.5 were used to identify transcriptionally distinct cellular clusters. For annotation, the most variable markers of each cluster were computed with *FindAllMarkers* setting to only return positives with a minimum of 0.25% and a logFC threshold of 0.5. Cluster annotations were performed based on DEGs and manually identified based on canonical markers and literature gene sets (**Table S4**)^40,41^. Composition analysis was performed to vizualize differences in cluster abundances between conditions using the Sccomp package (v1.0.0)^42^. Differences in gene expression between conditions was also assessed, and the DEGs were plotted in a volcano plot using *ggplot* (v 2_3.3.6)^43^. The expression of different transcriptomic signatures was assessed to characterize the differences among patients, timepoints, conditions and cell types. UCell suite^44^ was used to assign a score to each gene set (**Table S4**). Dotplot, violin plot and feature plot representations from the Seurat package were used to represent expression differences.

### Trajectory analysis

Trajectory analyses were performed with the Monocle 2 package (v2.24.1)^45^ to order cells in pseudotime, used as a measure of progress through biological processes based on transcriptional similarities. With this goal, we performed two studies: one including all T cells and the other with only CAR^pos^ T cells. The analyses started by creating “*CellDataSet*” objects using a negative binomial model. Then, the monocle algorithm was run using the 3000 most highly variable genes selected with Seurat (v4.3.0) after DDRTree dimension reduction and cell ordering. To plot the minimum spanning tree on the cells, *plot_cell_trajectory* function was used to visualize the ordered cells in the trajectory.

### Sc-αβTCR repertoire analysis

Sc-αβTCR repertoire sequencing data was analyzed using the scRepertoire package (v.1.7.1)^46^. The filtered Cell Ranger annotations were read in and TCR αβ chains were combined for all T cells based on their cell barcodes using the combine TCR function. Additionally, the *combineExpression* function was used for the integration of the combined αβTCR data with the scRNAseq data. Clonotypes were categorized by the absolute number (x) of T cells expressing individual clonotype sequences (the clonosize) as single (x=1 clonotype), small (1<x ≤ 5 clonotypes), medium (5<x<=20 clonotypes), large (20<x ≤ 100 clonotypes) and hyperexpanded (x>100 clonotypes), thus reflecting the actual clonal expansion for each sample. Occupied TCR repertoire space was calculated for each CAR^pos^ T cell subtype by counting the number of cells in each of the mentioned expansion categories. To assess the quality and quantity of the TCR repertoire of every CAR^pos^ sample, the diversity, richness and evenness were measured using *clonalDiversity* function by three different metrics: Shannon, Abundance-based Coverage Estimator (ACE) and the inverse Pielou measure of species evenness, respectively^47^.

To evaluate the expansion and ongoing proliferation of the CAR^pos^ T cell subtypes, a proliferation and expansion index was calculated. The hallmark genes involved in the G2/M checkpoint (GSEA, misigdb^48^) were used to calculate the proliferation index with *AddModuleScore* function from Seurat^36^. We also used startrac software to calculate the expansion index^49^. At the same time, the Morisita index was estimated by performing the overlap coefficient method to measure the similarity between the clonotypes from the different samples. The circlize package was used to visualize the interconnection of shared expanded clonotypes of the different T cell clusters^50^. Trend lines were drawn to visualize the scatter plots for expanded clonotypes shared in the IP and Peak^51^.

### Statistical analysis

Quantification and statistical analysis All statistical analyses were performed using R (v4.0.5). Statistical significance in each case was calculated using, two-sided Chi-squared test or Fisher’s exact test. To assess the association of CD4:CD8 ratio with EFS, we estimated a Cox’s proportional hazards model. The Kaplan-Meier survival curves were generated with survival R (v.4.3.0) package (v3.5-5). Where not indicated on the text, differences were not statistically significant.

### Data and code availability

Raw scRNAseq and scTCRseq sequences have been deposited in the Gene Expression Omnibus database under GSE235760 accession number. Processed single-cell data is available for interactive exploration in https://cellxgene.cziscience.com/collections/14dc301f-d4fb-4743-a590-aa88d5f1df1a. The code used to generate this analysis and images are deposited in https://github.com/Merguerrero/scCARTcells and https://github.com/mereulab/CAR-T_Figures. All other supporting data are available upon request.

## Acknowledgments

We thank CERCA/Generalitat de Catalunya and Fundació Josep Carreras-Obra Social la Caixa for core support. Financial support for this work was obtained from the Spanish Ministry of Economy and Competitiveness/European Union NextGenerationEU (PID2022-142966OB-I00, CPP2021-008508 and CPP2022-009759), the Horizon-EIC-2022-Transition (101113067), the ISCIII-RICORS-TERAV within the Next Generation EU program (plan de recuperación, transformación y resiliencia) and the Spanish Association against Cancer (PRYGN234975MEN) to PM; and the Health Institute Carlos III (ISCIII/FEDER, PI20/00822), Spanish Association against Cancer (PRYGN211192BUEN) and the Fundación Uno entre Cienmil to CB.

## Author contributions

MGM, ARH, JLT, DM, EM, MJ, AO, LMM, and CB performed experiments and interpreted data. AB, VOM, EAG, DB, NM, MJ, JD, AUI, AMS and AO led the clinical work, patient inclusion and follow-up for varni-cel, axi-cel and lisa-cel treatments, and provided clinical data. HH, EM, CB and PM supervised research and contributed key knowledge, techniques, and reagents. CB and PM conceived the study and funded the research. All authors actively participated in the manuscript preparation, with specific contributions from MGM, ARH, JLT, EM, CB and PM. All authors have read and agreed to publish the manuscript.

---

**Figure S1. Single-cell data exploration and quality**. **A)** Patient monitoring for the absolute number of CAR^pos^ T cells per µL over the first four weeks post-infusion. The red dots correspond to the peak of expansion. **B)** Violin plots representing expression level of the CAR sequence in CAR^pos^ and CAR^neg^ cells both for IP and Peak. **C)** Violin plots of the CAR expression by patient. Violin plots indicating the quality control metrics of T cell populations. Upper left panel, mean genes per cell; upper right panel, total number of reads; lower left panel, percentage of mitochondrial genes expressed; lower right panel, percentage of ribosomal genes detected. **E)** Dot plot showing the expression of five different gene signatures extracted from curated literature of T cell subtypes (Naïve, Stem Cell Memory, Memory like, Effector Memory, and Effector) in the identified T cell subpopulations. The Ucell method was used to score each signature (see Methods). **F)** Pairwise Morisita index, a statistic that represents the similarity between two populations, showing overlaps among all patient conditions (decimal points are indicated only inside circles; the higher the index corresponds to a more similar populations).

**Figure S2. Proportional and signature analysis in CAR^pos^ and CAR^neg^ T cells in the IP. A)** UMAP of the 38,190 cells pooled from all the samples from the 5 patients colored by the prediction of CD4/CD8 T cells identified by the well-established reference-based classifier ProjecTILs using a published dataset as a reference and Azimuth (see Methods). **B,C)** Activation (**B**) and tonic signaling (**C**) signature half violin plot of CAR^pos^ and CAR^neg^ cells within CD4 and CD8 predicted T cells. **D)** Scatter plot showing the percentage of CAR^pos^ CD4^+^ and CAR^pos^ CD8^+^ T cells at the IP in patients who were in CR (empty) or progressed (black) 2-years after treatment. Dot plots showing the expression of four literature-based exhaustion signatures in CAR^pos^ and CAR^neg^ T cells from each individual patient.

**Figure S3. Additional longitudinal analysis of CAR^pos^ T cells across timepoints**. **A)** Left panel, proportion of proliferative and non-proliferative CAR^pos^ T cells in the IP and Peak. Right panel, stacked bar plots showing proliferative clusters in the IP and Peak and for each patient. Cell populations colored as shown in Fig 2A. **B)** Boxplots showing the differential abundance analysis of CAR^pos^ T cell clusters in the IP and Peak. Expected abundance is indicated in grey boxes and the observed abundance colored by T cell subtype/cluster. **C)** Paired dot plots showing the abundance-based Coverage Estimator and Inverse Pielou scores in each patient in the IP and Peak to evaluate the diversity of the TCR repertoire within the CAR^pos^ T cells. **D)** Pseudotime analysis of CAR^pos^ T cells ordered by predicted pseudotime for each T cell subtype colored by timepoint. **E)** Total number of unique CAR^pos^ T cells clonotypes at IP and Peak per patient. **F)** Chord diagrams showing the interconnection of shared expanded clonotypes of the different CAR^pos^ T cell subtypes by patient.

