## Supplementary figures and images for "Integrative single-cell multi-omics of CD19-CAR^pos^ and CAR^neg^ T cells suggest drivers of immunotherapy response in B-cell neoplasias"

### Fig S1

**Guerrero-Murillo et al, Fig Suppl 1**

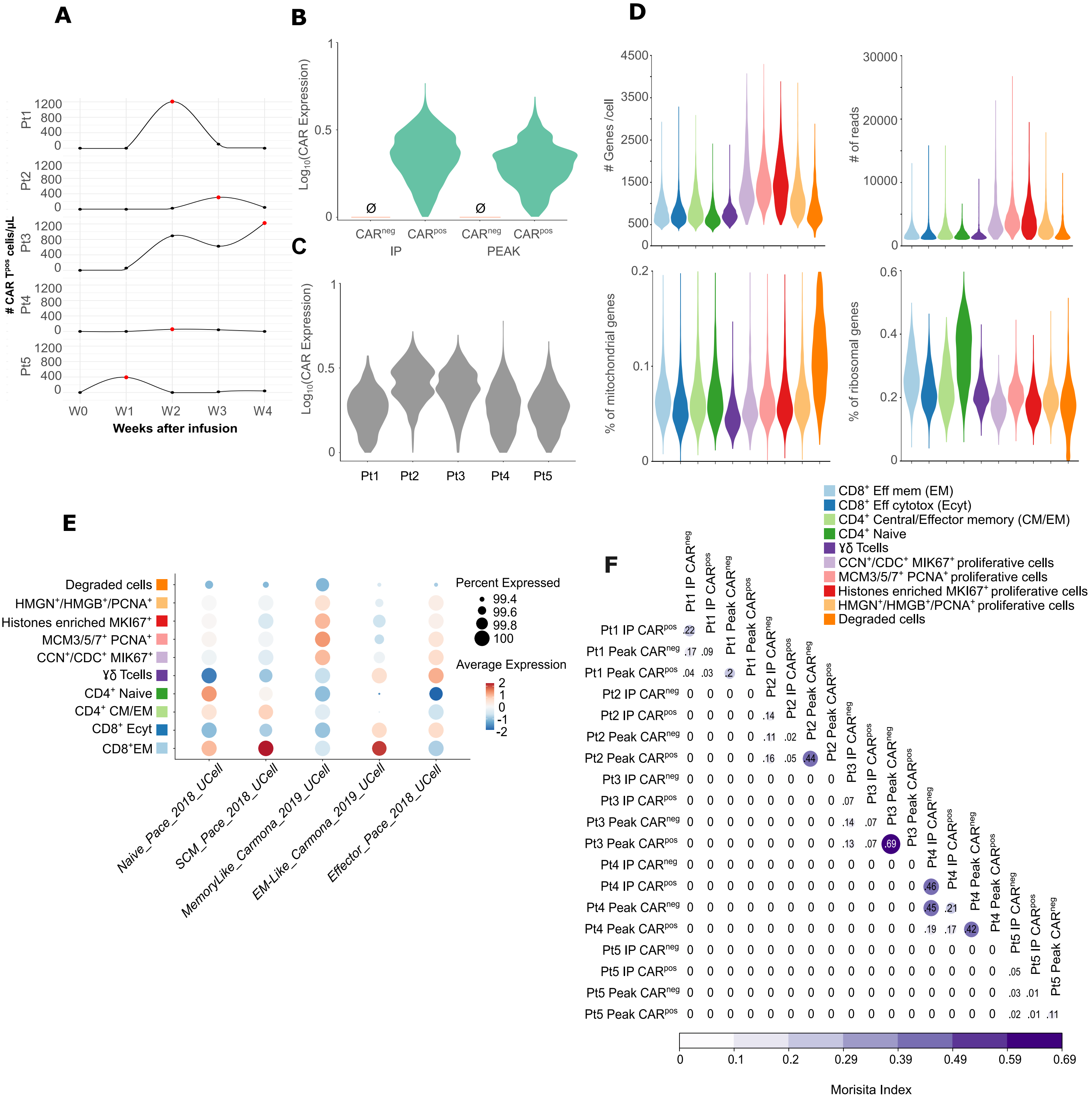

### Fig S2

**Guerrero-Murillo et al, Fig Suppl 2**

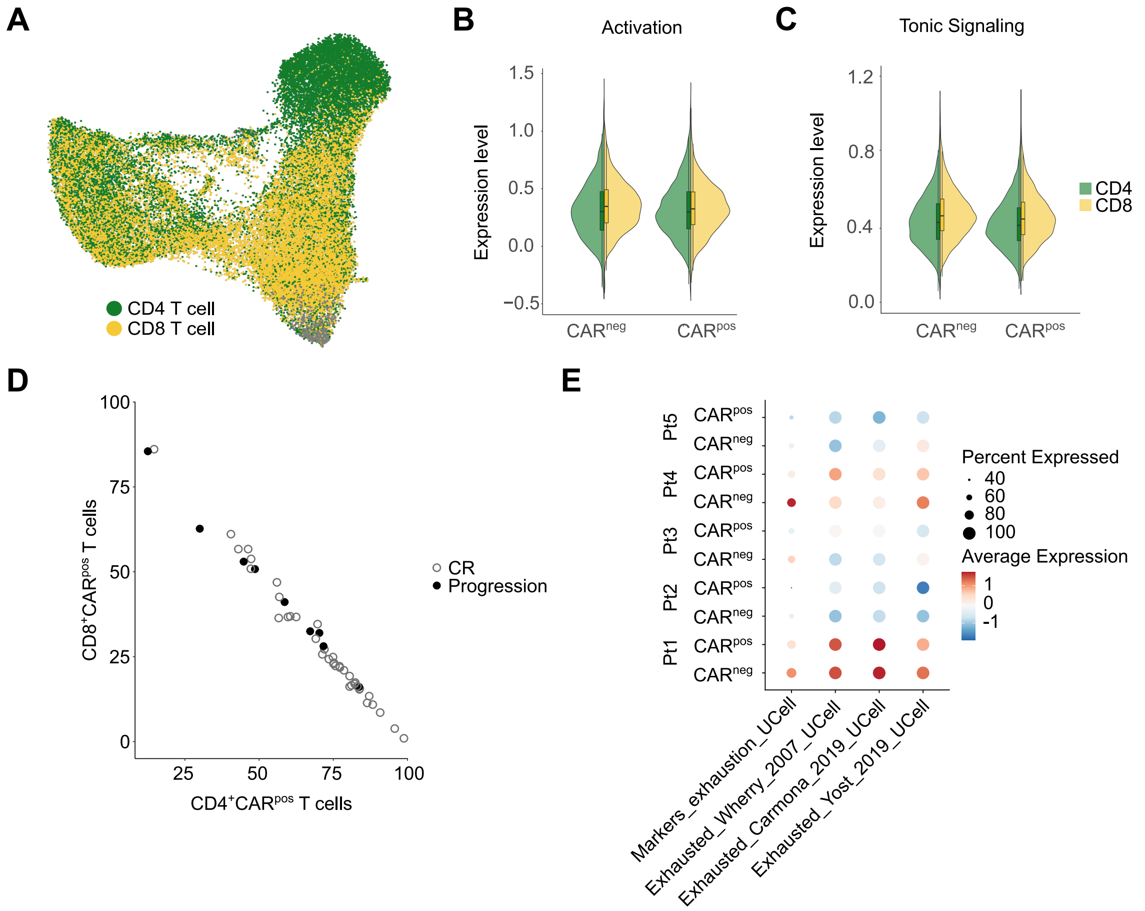

### Fig S3

Guerrero-Murillo et al, Fig Suppl 3

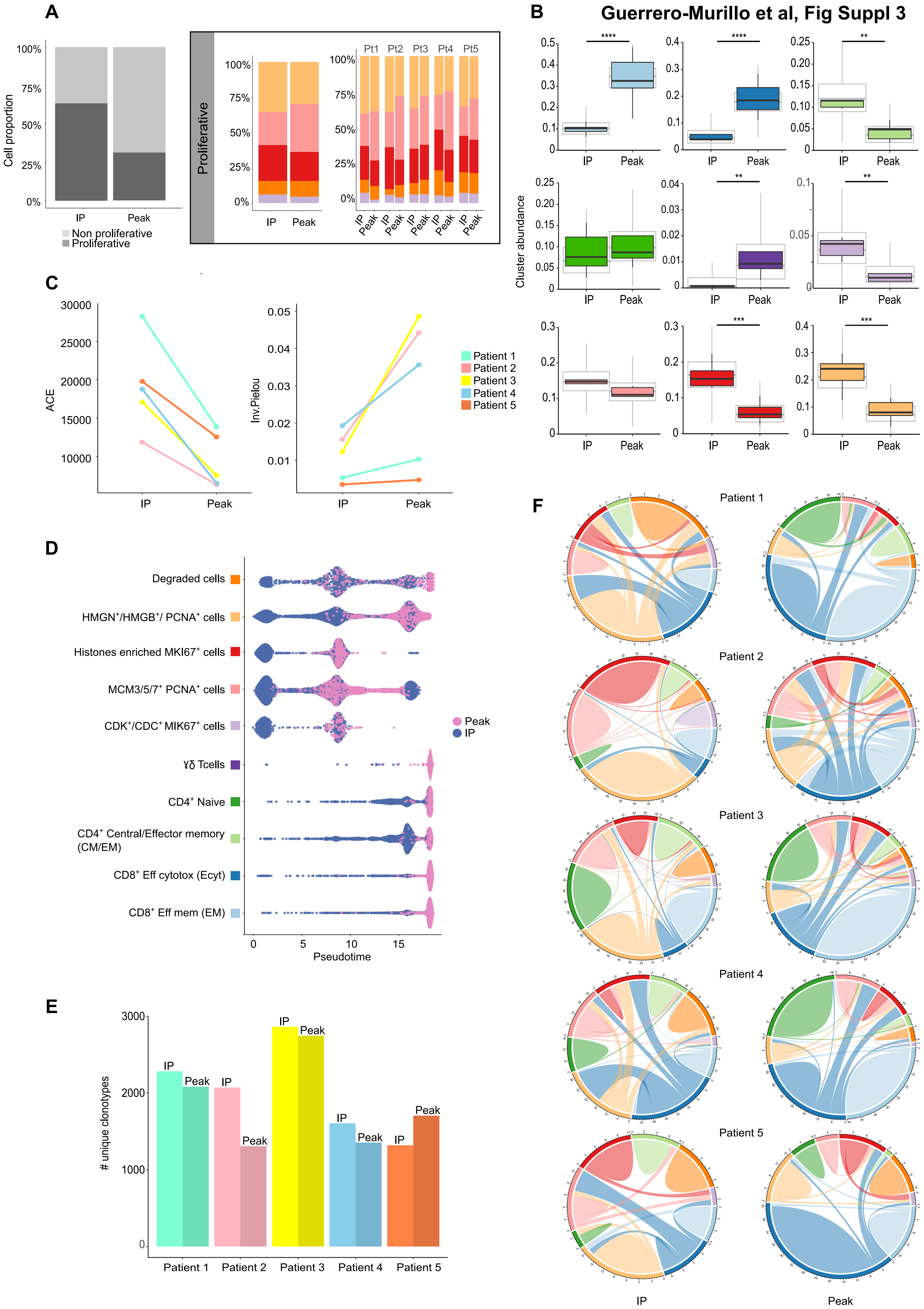
